# A Sparse Unreliable Distributed Code Underlies the Limits of Behavioral Discrimination

**DOI:** 10.1101/424713

**Authors:** Balaji Sriram, Alberto Cruz-Martin, Lillian Li, Pamela Reinagel, Anirvan Ghosh

## Abstract

The cortical code that underlies perception must enable subjects to perceive the world at timescales relevant for behavior. We find that mice can integrate visual stimuli very quickly (<100 ms) to reach plateau performance in an orientation discrimination task. To define features of cortical activity that underlie performance at these timescales, we measured single unit responses in the mouse visual cortex at timescales relevant to this task. In contrast to high contrast stimuli of longer duration, which elicit reliable activity in individual neurons, stimuli at the threshold of perception elicit extremely sparse and unreliable responses in V1 such that the activity of individual neurons do not reliably report orientation. Integrating information across neurons, however, quickly improves performance. Using a linear decoding model, we estimate that integrating information over 50-100 neurons is sufficient to account for behavioral performance. Thus, at the limits of perception the visual system is able to integrate information across a relatively small number of highly unreliable single units to generate reliable behavior.

## Introduction

Animals regularly identify the presence of external stimuli, and make decisions based on this evidence within very short time intervals^1-4^. Reliable performance with limited information requires a robust representation of the external world, but the structure of neural activity that underlies representation of sensory stimuli in circumstances where evidence is fleeting or scarce is not known.

In primates and carnivores, a natural timescale exists for integrating visual information - the fixation duration. Within an inter-saccadic duration (150-350 ms^5, 6^ cf. ~1000 ms for rodents^7^), the subject fixates on a part of the visual scene and extracts stimulus information relevant to behavior (orientation, motion, color etc.) suggesting that behaviorally relevant information can be extracted in a few hundred milliseconds. Rapid processing of sensory information has obvious evolutionary benefit^1, 4^, but the relationship between performance and neural representation has not been carefully investigated. One reasonable hypothesis would be that animals integrate information for a duration that leads to reliable responses in cortical neurons. We sought to address this possibility by carefully comparing the reliability of cortical representation with the quality of performance.

In several species^8-11^, the activity of neurons in the primary visual cortex (V1) enables conscious perception of visual patterns in the world (though unconscious blind sight effects do not require V1^8, 12, 13^). Within cortical visual pathways, visual information is thought to be represented by a sparse and distributed neural code^14, 15^. Such representations can be energetically favorable^16, 17^, have higher capacities than local codes, and are capable of generalization and tolerant to error. Constraining neural responses to be sparse and distributed in network models has recreated many of the properties of neurons in the early visual system^18^. Most stimulus parameters associated with evoking sparse responses nonetheless produce reliable stimulus-locked responses. It is not known if this response reliability is required for the system to extract meaningful information. Several additional questions remain unanswered: (1) How sparse can the responses be and still have animals perform reliable discrimination? (2) What “code” do animals use to detect and discriminate between stimuli? and (3) How many neurons are required to perform these discriminations? Answers to these questions have the potential to yield insight into how animals learn, integrate and process information over short timescales to support decision making.

In this study, we first establish that mice can rapidly integrate evidence over time to support decision making. Remarkably, we find that mice achieve plateau performance at timescales less than 100 ms. We then measure the electrophysiological responses of neurons across the layers of V1 to such short stimuli. These layers send and receive inputs to various other cortical and sub-cortical areas^19^ and could be involved in integrating relevant visual information. We find that there is only a marginal change in V1 neural activity under these conditions and the vast majority of neurons show no stimulus evoked activity even for stimuli in their receptive fields. Characterized by a measure of population sparseness (fraction of simultaneously recorded neurons that produced at least one spike in a time window spanning the visually driven response) high contrast and long duration (200 ms) stimuli engaged only 8% more neurons compared with no contrast. Furthermore, we find that L2/3 responses can be extremely unreliable (>85% of sensitive neurons failing to fire on a given trial). L5 neurons while more reliable still fail to fire on many trials.

To quantify how well individual neurons perform in discriminating visual stimuli, we developed a simple logistic regression model to quantify the contribution of each neuron to the discrimination task. We find that the vast majority of recorded neurons were poor discriminators with only a small fraction (~13%) consistently discriminating the orientation of the stimulus above chance. While reliable, these neurons never improved discrimination of the visual stimulus beyond a few percentage above chance. Based on the model, we can project the population requirement for the orientation discrimination task. We find that mice would need to integrate from a few tens to a few hundred neurons from individually unreliable responses to account for the reliable performance of mice in the orientation discrimination task. This constitutes a small fraction (<0.1%) of the total number of neurons available to encode the stimulus in V1 of the mouse^20^ indicating that mice can use sparse and highly unreliable neural responses to efficiently extract information to enable decision making at the limits of sensory perception.

## Results

### Mice performing orientation discrimination task integrate information over very short timescales

First, we trained adult mice in an orientation discrimination task. Naïve mice were introduced into the training arena with three response ports and their behavior slowly shaped to the appropriate response contingency (Figure 1A; see also Methods – Behavioral Training and Task Sequence and Parameters of Stimuli for Behavior). We captured the contrast dependence of the Orientation discrimination task in a series of trials where we varied the contrast of the discriminandum. As expected, we find that all subjects improved performance with increasing contrast (Figure 1B, grey lines, performance at c=1.0 > performance at c=0.15 for 8/8 mice). We fit logistic curves to the performance of subjects across contrast (Figure 1B, 1C, see methods Psychometric data fitting) and measured the threshold contrast (‘υ’ in Figure 1B, inset) for individual subjects to be between 0.06 and 0.30 (data not shown). Furthermore, the width of the tuning curve (ω, Figure 1B, inset) ranged between 0.11 and 0.50 (data not shown). To obtain population averages, we fit logistic regression curves on a simulated average subject (Figure 1B, blue curve, see methods Animal Variability and use of Average Subject). The average subject had a threshold contrast of 0.19, a tuning width of 0.48 with a lapse rate of 0.15 (Figure 1B). Most subjects perform OD with a threshold contrast around 15% and have plateau performance beyond 60% contrast. Even at the highest contrasts probed, performance was lower than 100% for each of the individual animals as well as the average animal (Figure 1B, blue curve). Thus, behavior of the mice shows a strong contrast dependence with performance plateauing around 80%.

**Figure 1.**
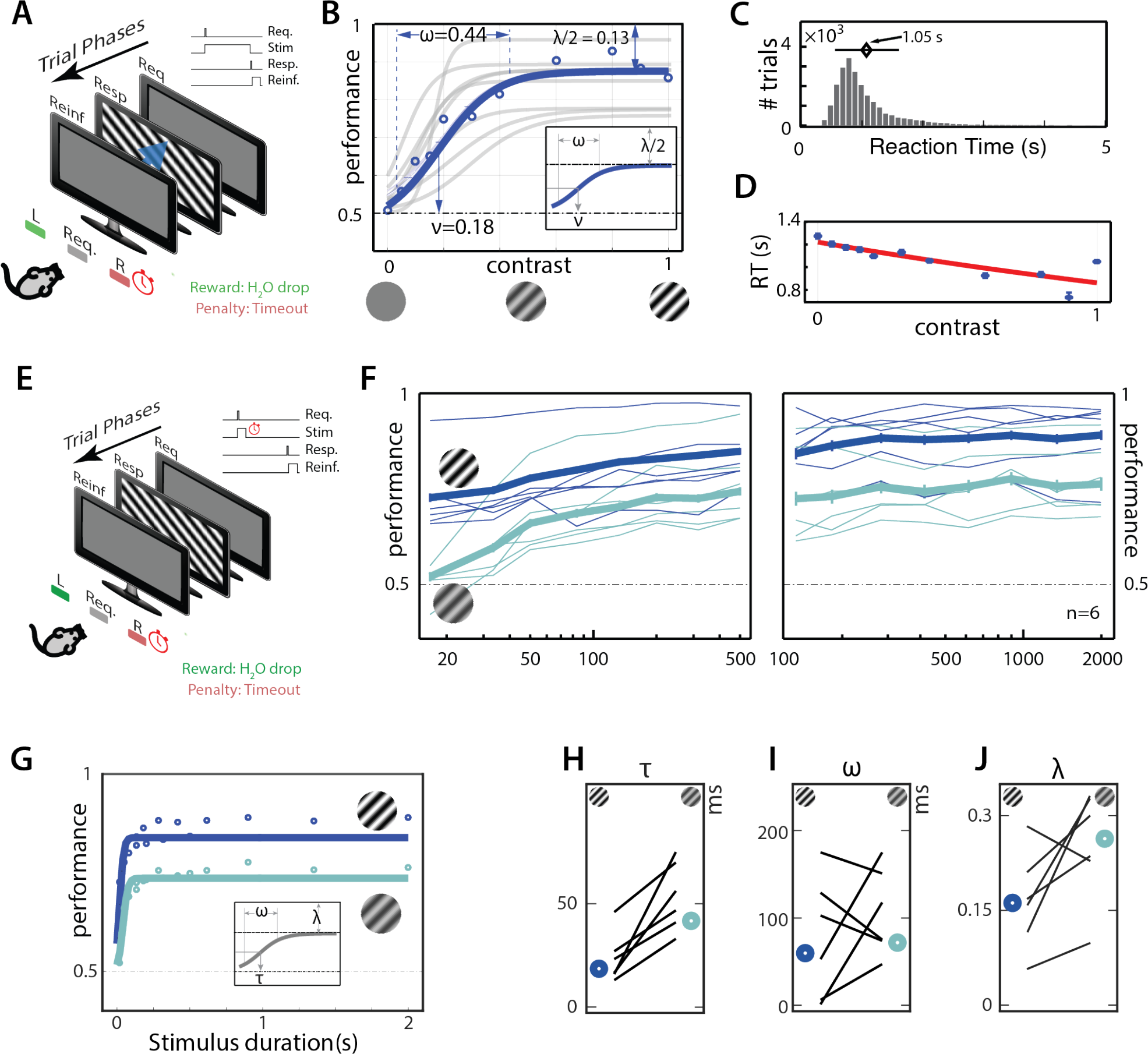
Mice integrate visual informatio*n*. **(A)** Schematic of trial structure *for contrast dependence of orientation discrimination*. **(B)** Performance improves with contrast. Sigmoid fits to performance of individual subjects (grey) and performance at specific contrasts along fits for the average mouse (blue) are shown. Also shown are the MLE of contrast at half-max performance (υ), tuning curve width (ω) and lapse rates (λ). **(C)** Distribution of reaction times for the average mouse. Average reaction time (black diamond) with 1 SD shown. **(*D)*** Average reaction time as a function of contrast. Data show expected and 95 % CI of mean reaction times. Best exponential fit is shown in red. **(*E)*** Schematic of trial structure *measuring for integration time*. Subjects control stimulus onset, but stimulus offset is under experimental control. **(F)** Performance improves with stimulus duration. Performance of individual subjects (thin lines) at high contrast (100%, dark blue) and at low contrast (15%, light green) as well as performance with 95% CI of average subject (thick lines) at high contrast and low contrast (light green) are plotted. **(G)** Sigmoid fits to performance of average subject at high contrast (dark blue) and at low contrast (light green) are plotted. Inset shows features of the sigmoid fit: minimal integration time (τ), integration width (ω) and lapse rates (λ). Maximal Likelihood Estimate of **(H)** minimal integration time (τ), **(I)** integration width (ω), and **(*J)*** lapse rates (λ) for the individual subjects along with the average animal (bold circle) at high contrast (dark blue) and low contrast (light green) are shown. Error bars indicate 95% percentile ranges of fits.

We next assessed the influence of integration time on performance. In this task, the animal had control over onset as well as offset of stimulus. We measured the Reaction Time (RT, duration between trial request and response) as a measure of the maximum amount of time over which visual information could have influenced decision making. RTs were positively skewed and median RTs for each animal ranged between 0.7s and 1.1s averaging 0.97±0.13s (mean ± sd, not shown). The median reaction time of the average mouse was 0.95s with a 95% CI between 0.54 and 2.5 s (Figure 1C). Mean reaction time was contrast dependent with higher mean at lower contrast (RT_C=0.15_ = 1.16s) compared to mean at higher contrasts (RT_C=1.0_ = 1.05s) (Figure 1D). This difference was small (104 ms) but was highly significant (Mann-Whitney-Wilcoxon U test, p<10^−17^).

To measure the time window over which mice effectively integrate information, we tested subjects trained in orientation discrimination to perform trials where the maximum duration of a stimulus available for the subject is systematically controlled (see methods, Task sequence). After this maximum duration (denoted ‘stimulus duration’), the screen changes to a gray screen and awaits response from the subject (Figure 1E). This trial structure puts limits on how long subjects can integrate visual information – stimuli after the stimulus duration do not contain useful information to perform the task. All subjects improved performance as the stimulus duration increased (Figure 1F, thin lines, performance at 500 ms > 16 ms in all animals, p<0.05, Agresti-Caffo statistics). This was true for high contrast (Figure 1F, blue lines) as well as lower contrast stimuli (Figure 1F, green lines). For example, while no subject had performance significantly above chance for a 16 ms stimulus at low contrast (c = 0.15), all subjects perform significantly above chance for stimuli lasting 50 ms at low contrast. At higher contrast (c = 1), all subjects were significantly above chance for a stimulus duration of 16 ms, the shortest stimulus duration tested. To precisely measure the dynamics of integration, we fit a logistic regression curve to the stimulus duration vs. performance curve (see methods, Psychometric data fitting, Animal Variability and Use of Average Subject, Figure 1G). This allowed us to measure the threshold integration time (τ), the integration width (ω) and lapse rate (λ). Based on these fits, individual mice had a threshold integration time between 13 and 46 ms at high contrast (c=1) averaging 24±12 ms (mean±sd, Figure 1H) and a threshold integration time between 33 and 75 ms at low contrast (c = 0.15) averaging 54±16 ms (mean±sd, Figure 1I). Thus, subjects require very short stimulus durations – significantly shorter than the typical saccade duration – to perform significantly above chance in an orientation discrimination task. The average subject had a threshold integration time of 18 ms at high contrast (Figure 1H, blue circle) and 45 ms at low contrast (Figure 1H, green circle), an order of magnitude lower than the mean reaction times (~1s) from the reaction time task described earlier.

Our assessment of the integration width – the stimulus duration over which performance improved – indicated that it was between 16 and 161ms (Figure 1I) for high contrast stimuli and between 45 and 168ms (Figure 1I) for low contrast stimuli. The integration width for the average subject was 123 ms at high contrast (Figure 1I, blue circle) and 94 ms at low contrast (Figure 1I, green circle). These measurements indicate that subject performance plateaus beyond 117 ms (72 ms width + 45 ms threshold integration time) even at the lowest contrast. This duration is an order of magnitude smaller than mean reaction times. The lapse rates for high contrast stimuli (0.33 ± 0.15) (Figure 1J) was consistent with the lapse rates of subjects at high contrasts (0.31 ± 0.14) in the reaction time task (c.f. Figure 1C, p = 0.85, Mann-Whitney-Wilcoxon U test). Similarly, plateau performance for low contrast stimulus (0.74 ± 0.08, mean ± sd) is comparable to the performance of subjects for low contrast stimuli (0.67 ± 0.1, mean ± sd) in the reaction time task (data not shown, p = 0.18, Mann-Whitney-Wilcoxon U test). This indicates that subjects have integrated as much of the information from the visual stimulus as possible within ~100 ms to guide their behavior.

### Activity of V1 Neurons is required for Orientation Discrimination

To understand the neural basis for fast visual integration, we recorded from neurons across the layers of primary visual cortex (V1) in subjects running on a Styrofoam ball while passively viewing stimuli of various contrasts, durations and orientations (Figure 2A). Visual responses of neurons from 15 subjects across a total of 81 sessions were captured. Of these, 65 sessions (N=9 mice) were recorded in naïve animals that had no exposure to the behavioral arena while the remaining 16 (N=6 mice) were recorded in animals that had prior experience in the orientation discrimination task and had threshold performance in the simple orientation discrimination task (Step 4, Table 2). In naïve animals, a total of 618 single unit activities and 926 multi-unit activities were recorded (Figure 2B-C, also see methods Single unit identification). These units had firing rates that ranged from a minimum of 0.22 Hz to a maximum of 60.25 Hz (Figure 2E). Most neurons fired very few spikes over the course of the session (Figure 2E, mode at lowest firing rate). Firing rates were dependent on the depth of the recording (superficial (L2/3): depth<400µm, deep(L5/6): depth>400µm, KS-test; p<0.001 Figure 2F). Spike widths varied from 0.07 ms to 0.43 ms based on full width at half maximum (FWHM) (Figure 2G). Spike widths and firing rates were negatively correlated (Figure 2H) and this correlation was highly significant (p<10^−9^).

**Figure 2.**
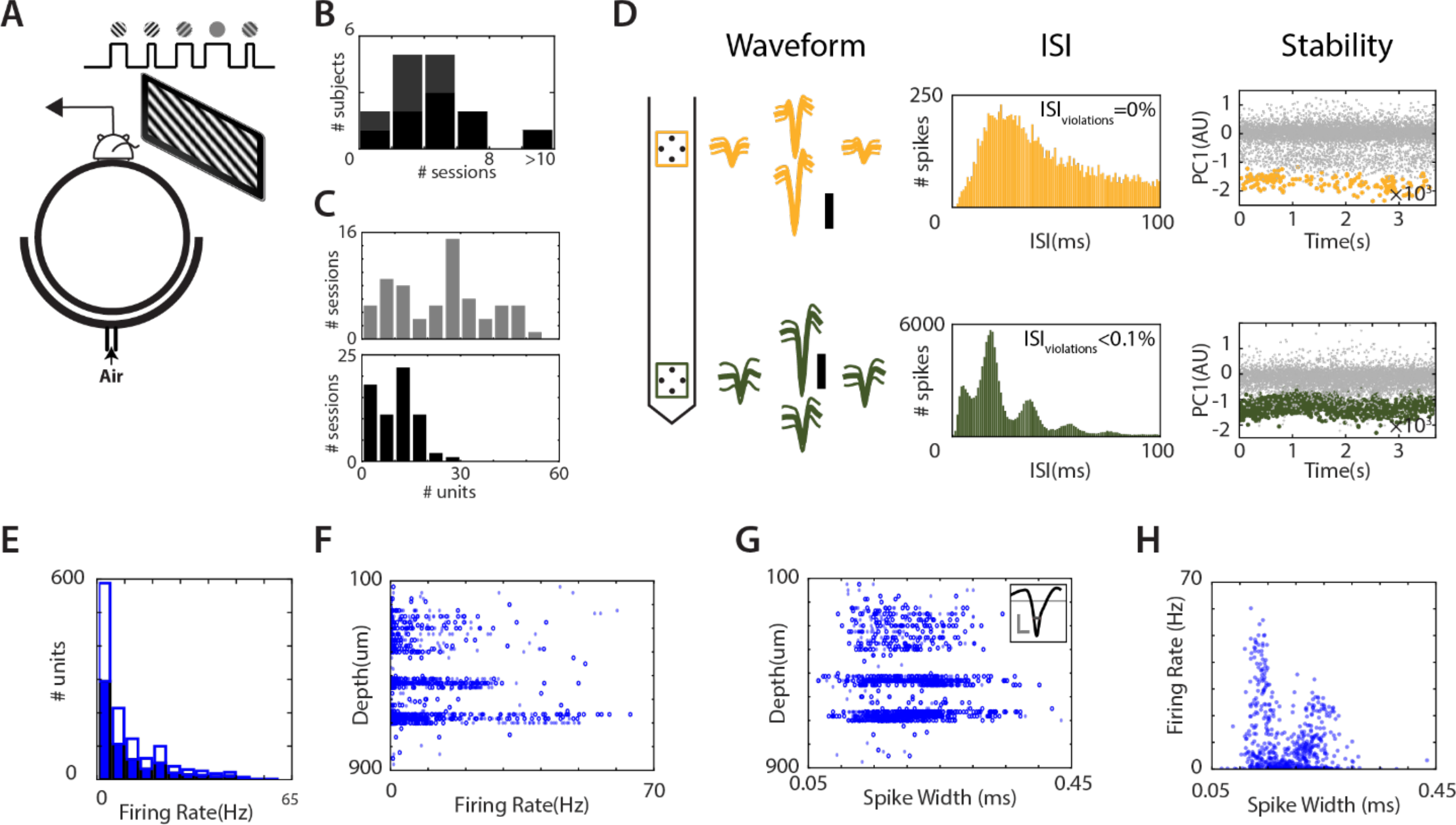
Statistics of Recording in V1. **(A)** Schematic of recording setup. (top panel) Subjects run on a Styrofoam ball suspended in air wile recording in the left primary visual cortex. We show visual stimulation to the right eye on an LCD monitor placed ~15 cm from the eye tangential to the eye. (bottom panel) Stimuli are gratings of different orientations, contrasts and durations separated by short periods (~1s) of gray screen. **(B)** Number of sessions collected from each subject (naïve: black; experienced: grey). **(C)** Distribution of total number of units (top panel) and well identified single units (bottom panel) collected in each session **(D)** Waveform, ISI, and first principal component of one single unit (top, yellow) and one multi-unit(bottom, green) simultaneously recorded in V1 on two separate tetrodes. **(E)** Distribution of firing rates of single units (solid) and multi unit activity (boxed) across all sessions. Firing rates **(F)** and spike widths **(G)** as a function of recorded depths. **(H)** Firing rates as a function of spike widths.

**Table 1.** Training steps for basic characterization of orientation discrimination.

| # | NAME | GOAL | STEP DETAILS | GRADUATION CRITERION | FIG. REF. |
| --- | --- | --- | --- | --- | --- |
| 1 | Free drinks | Learn to use reward ports | Reward ports occasionally squirt water | 2 days | NA |
| 2 | Earned drinks | „ | Mice need to lick different ports to receive water. | 3 days | NA |
| 3 | Luminance discrimination | Learn 2AFC trial structure | Large, high luminance stimulus appears on left or right side of monitor. Rewards only at side having high-luminance | 80% performance over the previous 200 trials | Supp. Figure 1b (blue trial blocks) |
| 4 | Optimal Orientation discrimination (OD-optimal)<sup>†</sup> | Learn OD on simple high contrast gratings | Circular patch of drifting gratings (2 Hz, 0.08 cpd, 100% contrast) tilted left or right (45°) to vertical. Rewarded in the direction of tilt. | 75% performance over the previous 200 trials | Supp Figure 1b (blue-green trial block) |
| 5 | Varied contrast<sup>†‡</sup> | Perform OD with varied contrast | Same as Step 4, except contrasts varied: $c \in [0,1]$ | Manual graduation | Supp Figure 1b (green trial block) |
| 6 | Varied Spatial frequency<sup>†</sup> | Perform OD with varied spatial freqs. | Same as Step 4, except spat. freqs. varied: $sf \in \{0.03, 0.06, 0.13, 0.25, 0.5, 1\}$ | Manual graduation | Supp Figure 1b (orange trial block) |
| 7 | Varied Orientation<sup>†‡</sup> | Perform OD with varied orientations | Same as step 4, except orientations varied: tilted $\theta \in [0, \pi/4]$ from vertical. | Manual graduation | Supp Figure 1b (yellow trial block) |

<sup>†</sup>: All Orientation discrimination stimuli had randomized orientation presented on every trial. The direction of drift was also randomized
<sup>‡</sup>: These steps contained completely ambiguous trials (0 contrast in step 5 and 0 in step 7). The correct response port was decided based on the nominal orientation of the stimuli (tilted right with 0 contrast or 0° gets rewarded in the right port).

**Table 2.**
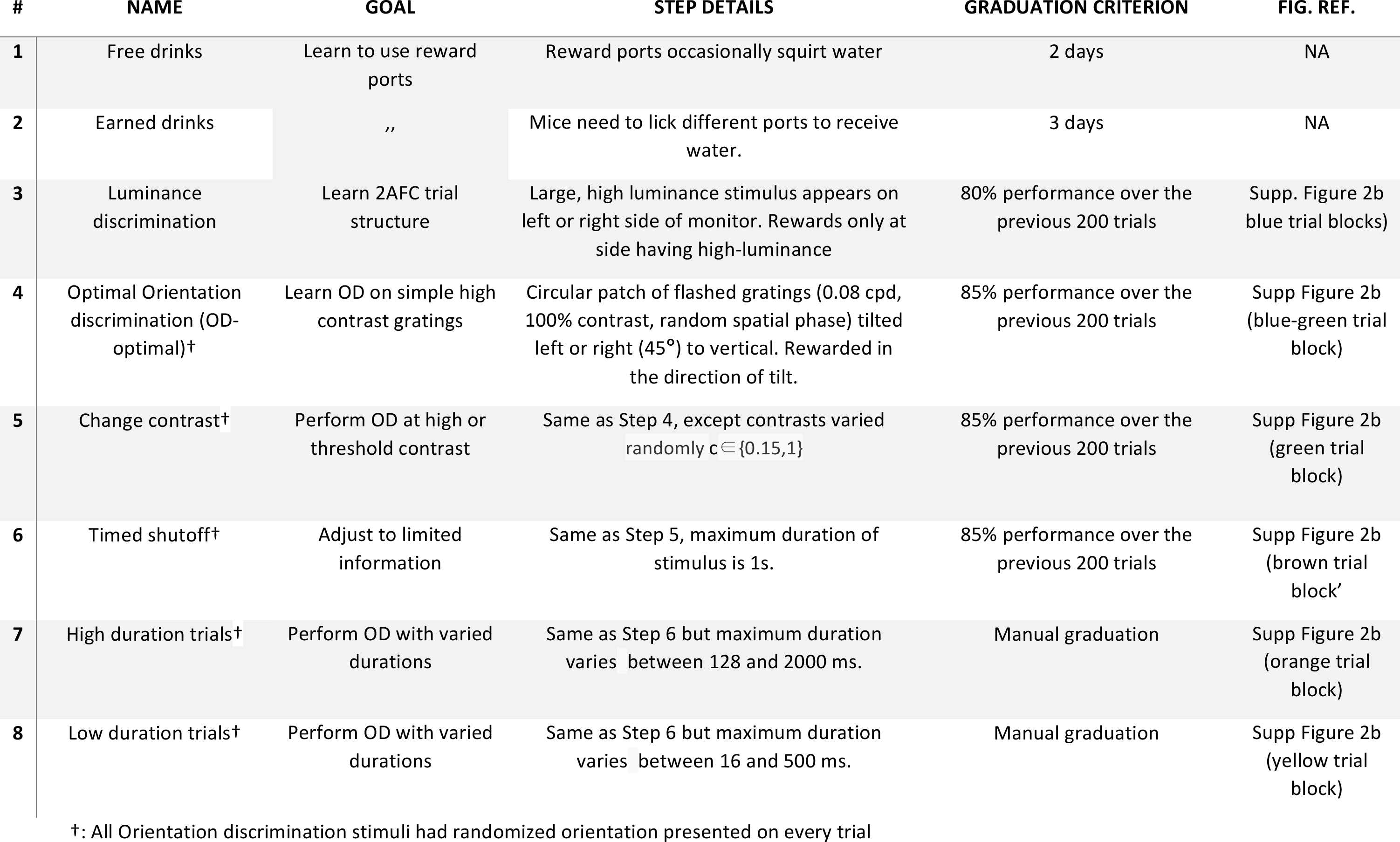
Training steps for measuring integration time.

| # | NAME | GOAL | STEP DETAILS | GRADUATION CRITERION | FIG. REF. |
| --- | --- | --- | --- | --- | --- |
| 1 | Free drinks | Learn to use reward ports | Reward ports occasionally squirt water | 2 days | NA |
| 2 | Earned drinks | „ | Mice need to lick different ports to receive water. | 3 days | NA |
| 3 | Luminance discrimination | Learn 2AFC trial structure | Large, high luminance stimulus appears on left or right side of monitor. Rewards only at side having high-luminance | 80% performance over the previous 200 trials | Supp. Figure 2b (blue trial blocks) |
| 4 | Optimal Orientation discrimination (OD-optimal)<sup>†</sup> | Learn OD on simple high contrast gratings | Circular patch of flashed gratings (0.08 cpd, 100% contrast, random spatial phase) tilted left or right (45°) to vertical. Rewarded in the direction of tilt. | 85% performance over the previous 200 trials | Supp Figure 2b (blue-green trial block) |
| 5 | Change contrast<sup>†</sup> | Perform OD at high or threshold contrast | Same as Step 4, except contrasts varied randomly $c \in \{0.15, 1\}$ | 85% performance over the previous 200 trials | Supp Figure 2b (green trial block) |
| 6 | Timed shutoff<sup>†</sup> | Adjust to limited information | Same as Step 5, maximum duration of stimulus is 1s. | 85% performance over the previous 200 trials | Supp Figure 2b (brown trial block) |
| 7 | High duration trials<sup>†</sup> | Perform OD with varied durations | Same as Step 6 but maximum duration varies between 128 and 2000 ms. | Manual graduation | Supp Figure 2b (orange trial block) |
| 8 | Low duration trials<sup>†</sup> | Perform OD with varied durations | Same as Step 6 but maximum duration varies between 16 and 500 ms. | Manual graduation | Supp Figure 2b (yellow trial block) |

<sup>†</sup>: All Orientation discrimination stimuli had randomized orientation presented on every trial

As expected from neurons in V1, a large fraction of neurons showed significant orientation tuning (Figure 3A-D). For a subset of neurons (389 single units and 722 multi units) we calculated orientation tuning (Figure 3B, polar plot) and vector sums of orientation tuning (Figure 3B, arrows) as measures of neural tuning to orientation (similar to prior techniques^21^) based on high contrast flashed gratings of different orientations that lasted 500 ms or drifting gratings that lasted 2000 ms. For all these neurons, we further calculated Jack-Knife error estimates of these measures by removing data from one trial at a time. Neurons showed varied selectivity and orientation preferences (Figure 3C, D). The orientation selectivity index spanned values from 0 to 1 with a mean selectivity of ~0.35, (Figure 3C) while the preferred orientation spanned the entire orientation space with significant preference for vertical and horizontal orientations (Figure 3D, p<10^−7^, KS-test vs uniform null hypothesis). We note that a small population of neurons (N=168) show very high orientation selectivity (OSI>0.95). These neurons have significantly lower firing rates compared to the rest of the population (0.21 Hz for High OSI vs 5.94 Hz for the population, p < 10^−68^, Mann-Whitney-Wilcoxon U test). The high OSI was due to extremely low firing rate for the orthogonal orientation, but each of these neurons had consistent orientation tuning – the standard deviation of Jack-Knife estimates of orientation preference was less than 20° in all high-OSI units with a median of 2.5°. Thus, as shown previously, we find that neurons in V1 are orientation tuned and are ideally poised to encode the stimulus orientation to direct the animal’s behavior.

**Figure 3.**
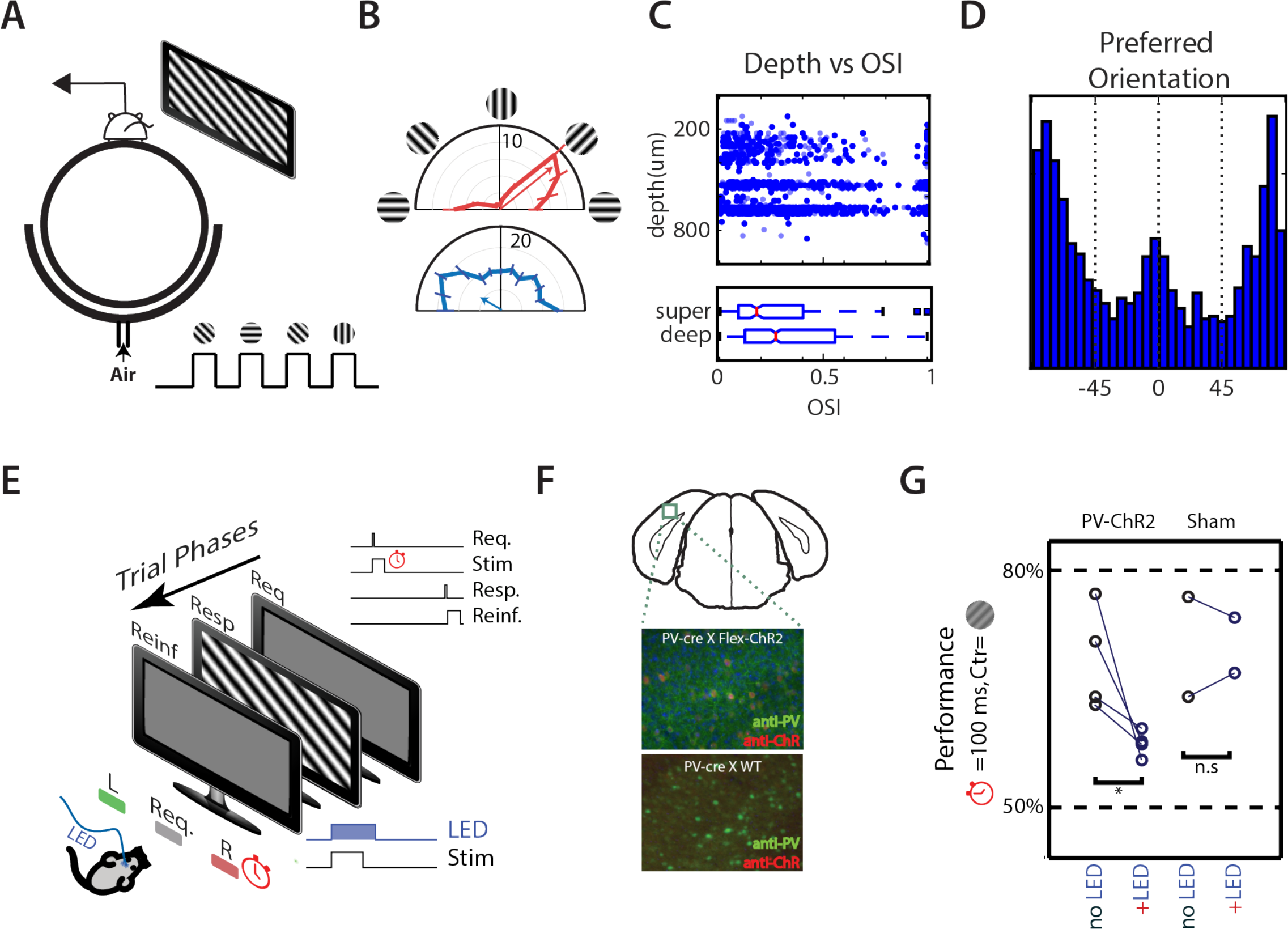
Neurons in V1 are orientation tuned and this activity is necessary for orientation discrimination. **(A)** Schematic of recording setup. (Top panel) Subjects run on a Styrofoam ball suspended in air while recording in the left primary visual cortex. We show visual stimulation to the right eye on an LCD monitor placed ~15 cm from the eye tangential to the eye. (Bottom Panel) Stimuli are gratings of different orientations separated by short periods (~1s) of gray screen. **(B)** Sample orientation tuning of two neurons with preferred orientation tilted to the right (top panel; red) and preferred orientation tilted to the left (bottom panel; blue). Arrows denote preferred orientation vector. Data are mean ±SEM of average firing rates. **(C)** Distributions of OSI across the population as a function of depth of the unit (Top panel; Solid blue - single units, light blue - multi unit) as well as box plots of the distributions for superficial(super.) and deep units (Bottom panel;mean OSI(superficial)=0.29<mean OSI(deep)=0.38, One-sided KS-test, p=3 × 10^−5^) **(D)** Distribution of preferred orientation calculated from the preferred orientation vectors. **(E)** Schematic of behavior. Subjects control stimulus onset, stimulus offset is under experimenter control. Stimuli last 100 ms and showed low contrast gratings (c=0.15). On random half of trials, a blue LED light is delivered to fiberoptic cannulae attached to the skull of the animal. **(F)** Epifluorescent image of visual cortical neurons with PV+ neurons in green and neurons expressing ChR2 in red imaged from coronal slices approximately over V1 (top panel) for mice expressing ChR2 (middle panel) and sham mice (bottom panel) **(G)** Average performance in trials with (‘+LED’) and without (‘no LED’) LED activation for four subjects expressing ChR2 in PV+ interneurons and two sham subjects not expressing Channel Rhodopsin. Average performance is significantly reduced for PV-ChR animals (paired t-test, p=0.0345) but not for sham animals (paired t-test, p>0.05)

To test if V1 neurons are required for orientation discrimination, we expressed blue-light sensitive Channelrhodopsin-2 (ChR2) in PV+ inhibitory interneurons (Figure 3F, top panel) in the primary visual cortex, which should suppress activity of projection neurons in V1 when activated. Suppression of cortical activity by the activation of PV+ interneurons is known to be immediate and reversible^22, 23^. We tested the causal role of V1 activity in behaving animals by activating Channel Rhodopsin on random trials while the subjects performed an orientation discrimination task (Figure 3E, also see methods Optogenetic Manipulation During Behavior). Performance in trials where ChR2 was activated was significantly lower than in trials where no ChR2 activation occurred (Figure 3G, ChR2 no LED vs. +LED; p=0.035, Mann-Whitney-Wilcoxon U test). We confirmed that the loss in performance was not due to the distracting influence of the blue light by performing identical experiments in sham animals that did not express ChR2 in PV+ inhibitory interneurons (Figure 3E, bottom panel). Performance in light activated trials was no different than the performance in trials without light activation (Figure 3G, Sham no LED vs. +LED; not significant, Mann-Whitney-Wilcoxon U test). Thus, activity of neurons in V1 is necessary for mice to carry out this orientation discrimination task.

### Visual responses in V1 are sparse

Next, we characterized the population responses of simultaneously recorded neurons to stimuli that last a short duration (<=200 ms). We recorded between 2 and 93 neurons simultaneously in our recording sessions. We show the stimulus (orientation, contrast and duration) (Figure 4A), spike counts in a time window that spanned the first 500 ms after stimulus onset of 25 (out of 39) simultaneously recorded neurons for the first 100 trials of one session (Figure 4B). This time interval (500 ms) was chosen for three reasons. First, previous reported latencies in mouse visual cortex indicate that for some of the stimuli tested, many neurons would not have begun their responses before the stimulus ended. Second, even if responsible neurons had begun responding to the stimulus, recurrent connectivity in cortex has the potential to drive activity for much longer than the stimulus duration^23^. And finally, preliminary analyses showed that for the stimulus set we used, a time interval of ~500 ms maximized the average information in the neural responses (data not shown). For each trial, the fraction of the recorded population that responded to the stimulus with at least one spike (Figure 4C) was computed. For the session shown, the average fraction of neurons that responded to the stimulus with at least one spike was 0.36 ± 0.09 (mean ± sd). Across all sessions and all trials, the fraction of L2/3 neurons that responded with at least one spike in the 500 ms window was 0.43 ± 0.17 (mean ± sd). This fraction of responsive neurons varied with the contrast and duration of the stimulus used (Figure 4D). We measured the mean population fraction for each session as a function of the stimulus parameters. For stimuli that lasted 100 ms, this mean fraction was significantly lower for zero contrast stimuli compared to the fraction at high contrast stimuli (Figure 4E, f = 0.37 at c = 0 vs f = 0.43 at c= 0.15, p = 0.037; and f = 0.37 at c = 0 vs f = 0.44 at c = 1, p = 0.015; Mann-Whitney-Wilcoxon U test). The difference between the fraction of responsive neurons was no different between stimuli of low vs stimuli of high contrast (p = 0.63, Mann-Whitney-Wilcoxon U test). On the other hand, the fraction of responsive neurons did not vary significantly with the duration of the stimulus (Figure 4F). These results indicate that for stimuli that drive reliable behavior, stimulus onset increases the fraction of responsive L2/3 neurons by at most 8%. This fraction was independent of the contrast or duration of the stimulus. Among L5/6 neurons, this fractional increase is greater (12%) but still represents a small minority of total neurons (data not shown).

**Figure 4.**
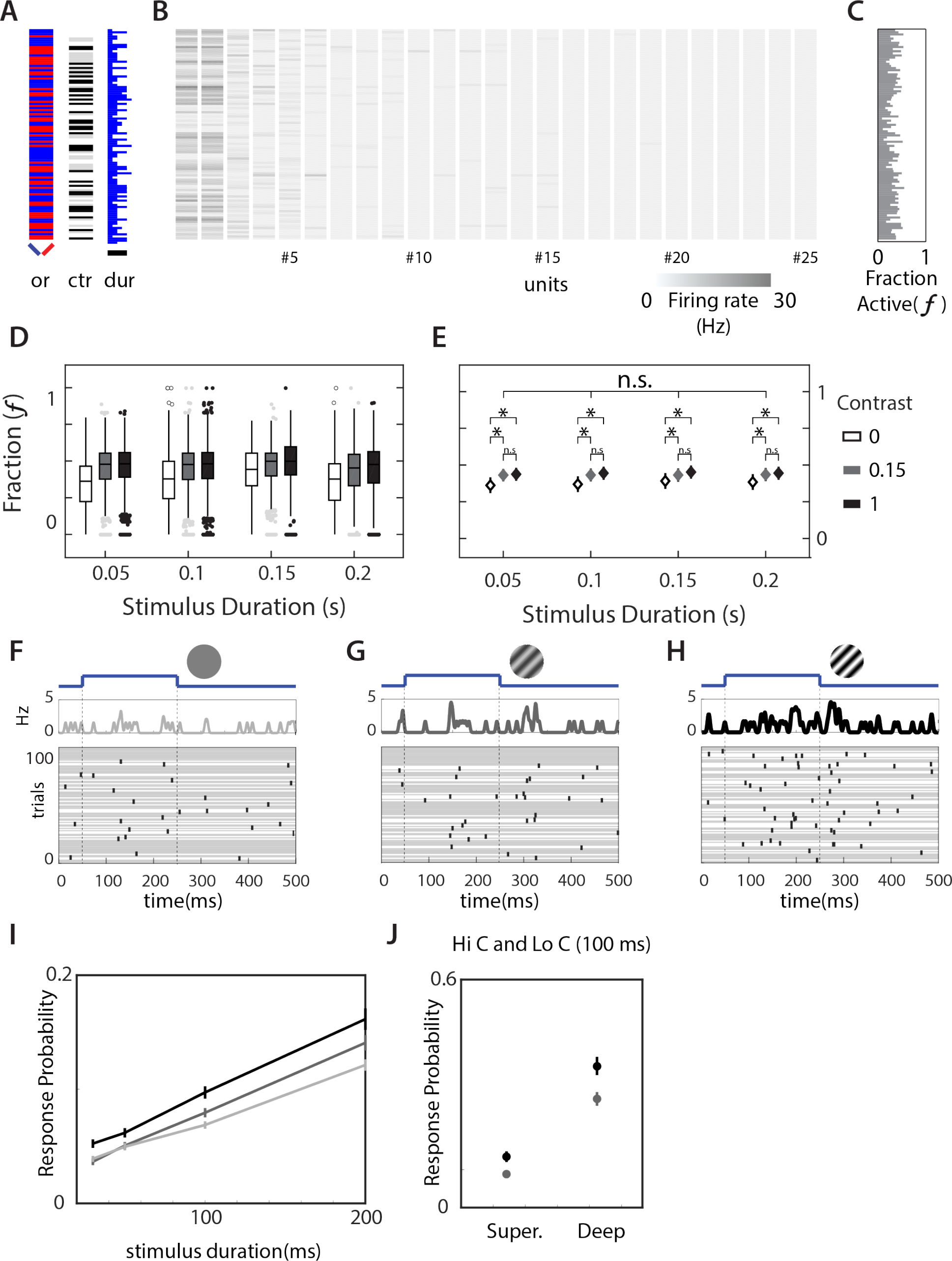
Population responses are sparse and unreliable. Schematic of a session. **(A)** The orientation (left = blue, right = red), contrast (contrast of 0 - white, 0.15 - gray, 1 - dark gray) and duration (black bar = 200 ms) of the stimulus of the first 100 trials of a single session. **(B)** Heat map of the spike rates of simultaneously recorded neurons in a time window that started with stimulus onset and ended 500 ms after stimulus onset. Inset shows the scale of the heatmap. This shows 25 out of 39 neurons recorded in this session. **(C)** The fraction of total recorded neurons that produced at least one spike in the time window of interest. **(D)** The distribution of fraction of responsive neurons for all trials pooled across all sessions plotted as a function of contrast and duration of stimulus. Box plots indicate medians and quartiles. Outliers are plotted separately. **(E)** Mean Fraction of responsive neurons across all trials for all sessions plotted as a function of contrast and duration of stimuli. Not all sessions contained stimuli for 0 contrast. Diamonds indicate mean across sessions. Error bars are 95% CI of mean (* indicates t-test, p<0.05, n.s. indicated t-test p>0.05). None of the comparisons across duration of stimuli for the same contrast was significantly different. **(F-H)** Raster of the responses of a single L2/3 neuron with preferred orientation to the right as a function of Contrast. Stimulus orientation and contrast (image) as well as onset and offset times (blue curve) are provided above the raster. In these panels, trials without a single spike are denoted by a gray line through the duration of the trial. **(I)** The probability of neurons responding to stimulus with a single spike at different contrasts(C-1, black; C=0.15, dark grey; C= 0, light grey) as a function of duration of stimulus.

### Visual responses in V1 are unreliable

Sparse responses could still underlie reliable behavior if individual neurons could respond reliably. To study the reliability of neural responses, we measured the probability that neurons would respond with at least one spike to stimulus presentation. We show the responses of one such L2/3 neuron with preferred orientation to the right of vertical (4F-H). The background activity of the V1 neurons was very low and they failed to fire in response to the visual stimulus on many trials, even at the highest contrast and duration (Figure 4H). To assess the influence of contrast on spike probability we calculated the fraction of trials which elicited at least one spike during the time that spanned the stimulus presentation and extended 100 ms after stimulus offset. The spike probability changed as we varied contrast at the preferred orientation (Figure 4F-H). Trials where the neuron failed to respond with a single spike are denoted as grey lines across the duration of the trial. The neuron responds unreliably even at the highest contrast (Figure 4H) and this reliability only reduces for lower contrasts (Figure 4F, 4G).

To measure the contrast and duration dependence of response probability, we split the recorded population of neurons into populations sensitive to orientation tilted to the right of vertical and to the left of vertical. Across the population, even at the highest contrast and highest duration presented, neurons fire spikes on fewer than 20% of trials (Figure 4J, black) on average. Responses at lower contrasts (Figure 4I, grey) and durations elicit spikes on fewer trials. Thus, for the stimulus conditions that drive reliable behavior (low contrast, 100 ms stimuli/high contrast 50 ms stimuli) neurons that are responsible for encoding that behavior fire fewer than 0.1 spike every trial.

### Individual neurons encode short stimuli poorly

Given the unreliability of individual neurons, we used the trial-to-trial responses to decode the orientation of stimuli. We split the responses of individual sessions across differing durations, orientations and contrasts into “training” (70%) and “test” (30%) sets (Figure 5A, See Methods, Fitting Performance of Individual Neurons and for Populations of Neurons in a Session). A logistic regression model was created on the training set and used on the test data to obtain decoding performance for the neuron (Figure 5A). To ensure that the choice of trials did not bias the performance of the logistic regression model, we fit the model to different random subsets of trials (splits) 100 times and used the average performance across these different subsets as a measure of the decoding performance of the neuron in predicting the stimulus. Individual neurons encoded the orientation of the stimulus poorly. Average performance across these 100 subsets across all stimulus conditions varied from 0.44 to 0.59 (Figure 5C), compared with chance performance of 0.50. The process of fitting a logistic regression to the responses did not always improve prediction, even for the training dataset. However, a small subset of the neurons (13.8%) showed reliably improved prediction in at least 70 out of 100 training sets. The performance of this subset of neurons, tested on the corresponding “test” dataset, (Figure 5C, Blue) was marginally but significantly higher than the overall population (0.52 vs 0.5, p < 10^−19^, Mann-Whitney-Wilcoxon U test). This subpopulation had units whose orientation preference was enriched in angles around the discriminated stimulus (data not shown). Specifically, this orientation preference distribution was significantly different than a flat distribution (KS-test, p<0.01) as well as the distribution of orientation preference of the overall population (KS-test, p<0.01). Within this sub-population, performance increased with contrasts and durations in a manner consistent with the improvement in performance of subjects in the orientation discrimination task (cf. Figure 5D vs Figure 1F). Nevertheless, even the best performing individual neuron was not close to the performance of the whole animal (e.g. ~80% behavioral performance for full contrast stimuli that lasted 100 ms or ~70% for low contrast stimulus that lasted 100 ms), indicating that any strategy that relies on the response of an individual high performing neuron was unlikely to drive the behavior of the animal.

**Figure 5.**
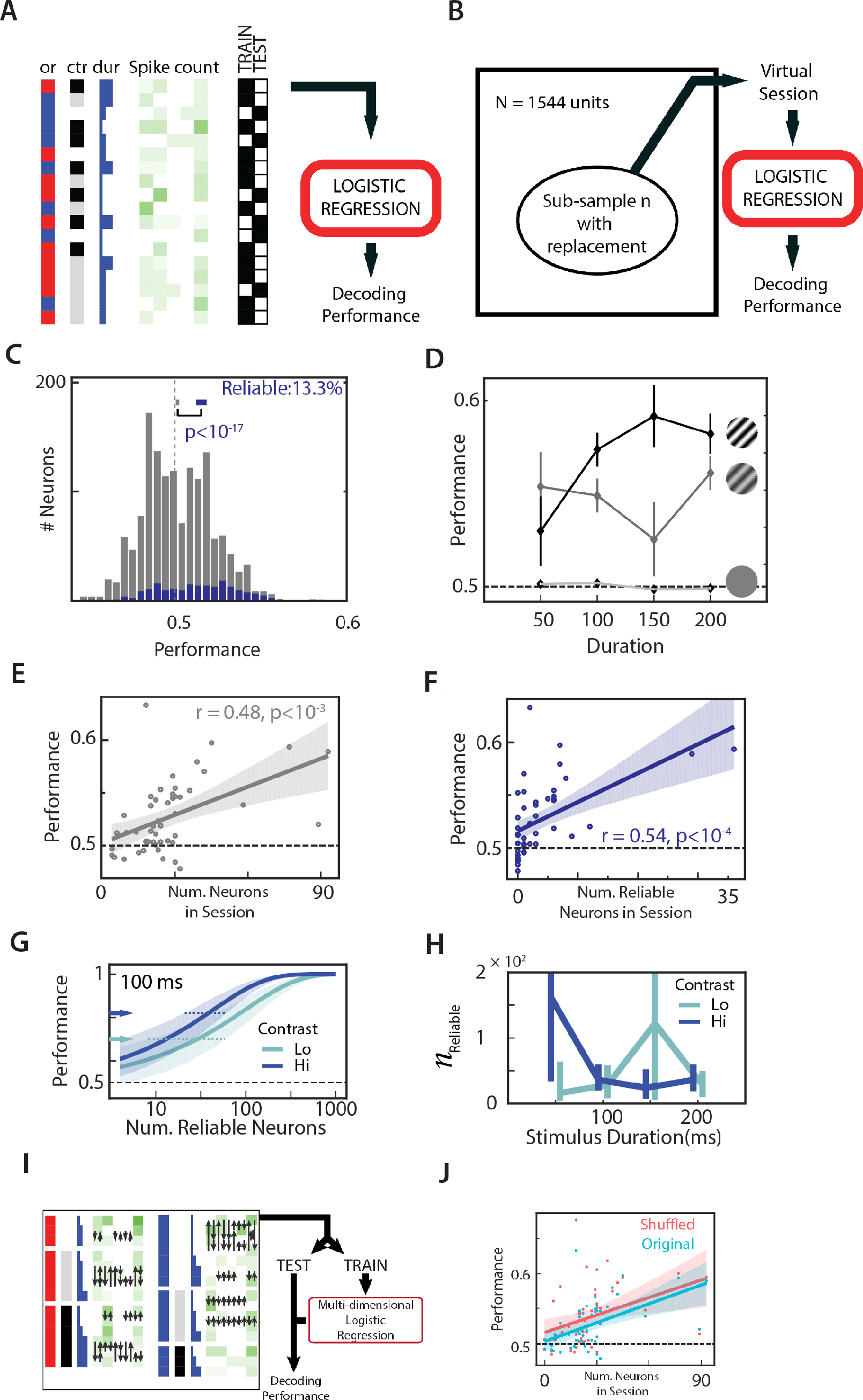
Integrating evidence across neurons improves performance. **(A)** Schematic of logistic regression fit. The responses of the neuron were split into 70% training (TRAIN) and 30% testing (TEST) and spike counts were used as features for a logistic regression. Regression coefficients calculated from the TRAIN dataset was tested on the TEST dataset to obtain a decoding performance. **(B)** Schematic of resampling method to create virtual session of different subpopulation of neurons. ‘Virtual’ neurons were subsampled with replacement across the entire dataset and spike counts of these neurons simulated by sampling from their respective response distribution. Prior logistic fits were then used to estimate performance **(C)** Histogram of decoding performance of neurons (calculated as in A, with individual spike counts as features) in the recorded population calculated as the average decoding performance across ten independent splits of training-testing subsets. A performance of 0.5 indicates no information. Neurons that reliably improve the logistic fit upon inclusion no matter how the data is split into training and testing is shown in blue. **(D)** Duration and Contrast dependence of performance of neurons. Data show mean ± 95 % CI performance for 0% (unfilled), 15% (grey) and 100% (black) contrasts. Multinomial logistic regression fits are obtained from training data and tested on test data as a function of **(E)** total number of neurons in the session or **(F)** total number of reliable neurons in the session. Circles indicate performance of individual sessions. Solid line indicates best linear fit and shaded area indicates the confidence interval of the linear fit. **(G)** Relationship between number of reliable neurons and performance measured for virtual sessions created as in D for gratings of high (blue) or low (green) contrast that lasted 100 ms. Solid lines indicate median performance and shaded areas indicates 95 percentiles of performances across 10000 sub-samples. Arrows indicate the measured behavior for low and high contrast stimuli lasting 100 ms and dashed lines indicate the number of neurons required to perform as well as the animal. **(H)** Number of reliable neurons required to perform as well as an average subject as a function of stimulus duration for high contrast (blue) and low contrasts (green). Data shows median ± 95 percentiles. **(I)** Schematic of shuffling strategy for estimation influence of correlations on performance. Stimuli were grouped according to stimulus parameters (orientation, contrast, duration). Spike number swaps (black arrows) occurred only between stimuli of identical parameters. **(J)** Same as in E but for Original (blue) and Shuffled (red) datasets.

### Integrating information across neurons predicts orientation

To determine how decoding of stimulus improves with integration of information from multiple neurons, we used the simultaneously recorded spike rates in all the neurons recorded in a session to fit a multinomial logistic regression similar to the method employed for the single neuron fits (see Methods, Fitting Performance of Individual Neurons and for Populations of Neurons in a Session). Performance with multinomial logistic regression was directly correlated both with the number of simultaneously recorded neurons in the session (Figure 5E, r=0.48, p<10^−3^) as well as with the number of reliable neurons recorded in the session (Figure 5F, r=0.54, p<10^−4^). There was rapid increase in performance with recruitment of additional neurons indicating a high sensitivity to performance of integrating information from even a relatively small number of neurons. To estimate the number of neurons required to perform as well as the animal, we simulated new sessions using increasing subpopulation of neurons from the overall population. Each neuron in the subpopulation was selected at random with replacement from the original population and firing rates for multiple trials simulated from known responses of the neuron for similar stimuli (Figure 5B, also see Methods, Simulating Neural Subpopulations and Measuring Performance of Simulated Populations). We used these virtual sessions along with known logistic fits for individual neurons to create a multinomial logistic regression (Figure 5B). This analysis revealed that performance improved with the number of neurons used and was dependent on the contrast (Figure 5G; 100 ms stimuli of high and low contrast) and duration (not shown) of the stimulus. We used the population dependence curves along with known performance of the average subject in the behavioral task to estimate how many reliable neurons would be required to decode the stimulus as well as the average subject could. Across stimulus conditions, a few tens to a few hundred neurons are required to match the performance of subjects from behavioral experiments (Figure 5H) averaging 49 neurons for low contrast stimuli and 65 neurons for high contrast stimuli. This still represented <0.1% of the neurons in V1 and suggests that information contained in the sparse and variable firing pattern in V1 is sufficient to account for visually-guided behavior. Our resampling analysis of the population dependence of performance removed all covariations in the responses of neurons. To estimate if such correlations were important for decoding, we created virtual sessions that removed any correlations present in the neural responses within a session by swapping spike numbers across trials with matched stimulus parameters (orientation, contrast, duration). We then fit multinomial logistic regressions to the shuffled sessions (Figure 5I) similar to the procedure described earlier. On average, shuffling improved the performance marginally but significantly (Δperformance = 0.012, p<0.001, Mann-Whitney-Wilcoxon U test, not shown). However, this did not significantly affect the slope of the Performance vs Number of neurons curve (shuffled, red vs. original, blue, Figure 5J) indicating that the correlations have little effect on the decodability of the stimulus. Thus, our population estimates are a reasonable approximation to the real requirements of animals discriminating gratings based on a spike rate code.

## Discussion

Our assessment of the timescale of evidence integration for visually guided decision making was enabled by the development of a high-throughput platform for studying visual function in rodents^24^. We adapted this platform for mice such that dozens of mice would perform hundreds of trials daily and quickly learned many kinds of visually directed behaviors. By simultaneously training and testing multiple subjects programmatically, we were able to collect behavioral information at scale, allowing us to measure psychophysical tuning curves by changing parameters of the stimulus used to probe the behavior^24^. We first measured the contrast tuning of orientation discrimination (N=8 mice). The threshold contrast measured (~15%) was similar to thresholds measured earlier in mice^25-28^, comparable to those of the hooded rat^29^ but is an order of magnitude higher than contrast thresholds in monkeys and humans^30^.

Nevertheless, mice improve performance with stimulus duration and thus are capable of evidence integration across time. This is similar to the behavioral responses in rats^31^, but potentially different from responses in monkeys and humans^32^.

One of the principal findings of this study is that mice can extract sufficient information on very short timescales (tens of ms) to perform visually driven tasks. Moreover the useful integration times are an order of magnitude shorter than typical stimulus duration used to probe visual perception in mice^26, 27, 33, 34^. The maximal integration time – time beyond which subjects do not integrate visual information – was ~200 ms. This may not reflect an inability of mice to integrate information over longer time durations and might just reflect the strategy subjects use to maximize reward given the non-zero cost of accumulating evidence ^35, 36^. Indeed, in other tasks rats are known to integrate information over many seconds^37^. Nevertheless, subjects that integrate information quickly perform well in the orientation discrimination task.

The ability of mice to discriminate orientation requires V1, as their performance is compromised by optogenetic inhibition of V1 responses. This is consistent with known response properties of V1 neurons^38, 39^ and with prior research in rodents^26^ investigating the requirement of various visual pathways for perception. At these short time scales, we find that responses in V1 are sparse and unreliable. We find that the average neurons can fail to produce even a single spike to preferred orientations on 85% of the trials on average while still allowing maximal performance (>85% correct) in subjects. The ability to discriminate orientation from responses of individual neurons based on a spike number code varied between 0.43 and 0.58 indicating that the best performing neurons do not come close to accurately predicting stimulus orientation. This stands in contrast with earlier work^40-42^ indicating that some V1 neurons) can be as sensitive or more sensitive than the subject. It remains possible that task engagement might increase the reliability of the neurons in such a manner that neurons become highly reliable. However, this increase in reliability needs to be large and prior estimates of reliability changes with training do not show such large increases^43^. To assess if training in the behavioral task improves the reliability of the neurons, we recorded from an additional cohort (N=6 mice, 16 sessions) that had experience with the orientation discrimination task. We found that the response probabilities measured from these animals were not significantly different from those measured in naïve animals. Recording in the presence of task engagement might further change these responses, however, we do not have data to address that question. Since the animal performs much better than would be predicted from single unit responses, we sought to explore the impact of pooling information over populations of neurons on stimulus prediction. Based on the approximation that firing rates are uncorrelated, we estimate that pooling across ~50 reliable neurons would be required to perform as well as the subject. Prior estimates of the population requirements of neural coding has indicated that very few neurons may be required for certain kinds of sensory discriminations in mammals^40, 41, 44, 45^. However, the minimal required population requirements measured using sparse optical activation to be comparable to that measured in this study^46^. The mouse V1 contains ~ 1 million neurons^20, 47, 48^ in each hemisphere. Thus, the orientation discrimination task requires only a small fraction of the overall population (<0.1%) for adequate performance.

We find that the performance of the animal peaks at about 80%. Given the number of neurons available to encode the stimulus, and the small fraction required to perform as well as the animal, why don’t subjects use more neurons to improve performance? Our estimate of the population requirement is based on a neural decoding model that uses only trial-by-trial spike counts. Models that use average firing rates within a time period to indicate the strength of the stimulus have a long history in the literature^49^ and are widely used^50-52^. However, cortical responses can be temporally precise under appropriate conditions^53, 54^. Many variations of a temporal codes have been proposed including models that consider latency to first spike^3^, phase of spike relative to underlying oscillations^55, 56^ or co-activation of ensembles of neuron^57-60^. However, this temporal precision typically comes with improved information flow within the brain^61-63^ and if used should reduce the population requirement below that measured in our data. Thus, such temporal codes cannot explain why subjects are unable to perform better.

While coordinated temporally precise firing patterns cannot explain performance in OD tasks, another form of coordination among neurons might affect decodability: Neural responses within cortex show significant correlation with one another^64, 65^. Covariations in neuronal responses prevent pooling responses across multiple neurons from removing noise^66^ and therefore might reduce the overall information available to the subject. Indeed, the effects of attention on improving perception are thought to largely be due to changes in the correlation^67, 68^. Thus, it is possible that correlations across V1 neurons limit the total information the animal can extract from the population. Alternatively, however, correlation could make decoding stimuli easier if it is another channel to communictae information about the stimulus present in the animal’s environment. In this situation, including correlations might improve the code^69, 70^. We investigated the effects of correlation by artificially removing them from our recorded sessions by shuffling neural responses across trials (Figure 5 I, J). While shuffling improved performance on average, it did not substantially change the slope of the Performance vs. number of Neurons curve (Figure 5J). This suggests that correlations play only a marginal role in affecting the cortical code underlying orientation discrimination at the limits of perception. However, even small correlations can affect the decoding of stimuli if information is spread across large populations of neurons. We have only tested the role that correlations play for population sizes between 1 and ~50 neurons. Based on the population requirement for reflecting behavioral performance (~50 reliable discriminators) and the measured prevalence of reliable discriminators (1 in 7.5), we would then need to measure the influence of correlation on simultaneously recorded population of size ~375 neurons. We have not considered the effects of correlations on such a large population in our models. Nevertheless, L2/3 neurons we record from are known to receive many inputs from nearby L2/3 neurons and are likely to exhibit significant correlations in responses^71, 72^. Future experiments should aim to record from such large populations and directly measure the effects of correlations on decodability.

Finally, given the unreliability of individual neurons and the evidence that behavioral performance is dependent on pooling information from a population of neurons, it is worth asking how such pooling would be achieved in the cortex. Both L2/3 neurons and L5/6 neurons are known to receive information from L2/3 neurons. So, these are likely populations whose firing would reflect pooled population response. The fact that even the best performing L2/3 neurons are poor predictors of performance, and previous data indicating L2/3 neurons have a bias towards making connections with other L2/3 neurons with similar Orientation Selectivity^73, 74^ suggests that L2/3 may not be the site of population pooling. L5/6 responses while more reliable, are still not enough to perform as well as the animal in such a behavioral discrimination task. This again eliminates L5/6 as the location of the population pooling. Thus, the site of population pooling must lie outside V1. Careful dissection of the visual pathway through optogenetic inactivation of secondary visual areas along with recording neural responses in these areas may be needed to definitively identify the integrator. Nevertheless, this integrator must be capable of producing reliable outputs from the unreliable outputs of V1. Our findings provide evidence that visually guided behavior at the limits of perception relies on effective integration of information across units with sparse and unreliable responses to stimuli.

## Methods

All procedures were performed with the approval and guidance of the Institutional Animal Care and Use Committee (IACUC) at the University of California, San Diego. We used N=35 adult male and female mice for this study.

### Behavioral Training

Behavioral training methods were adapted from training systems developed previously for rats ^24^. Water restricted adult (>P90) male and female mice were trained to use an operant conditioning chamber to receive water rewards while performing visually guided tasks. Water restriction and behavioral training/testing continued five days a week followed by weekends where subjects received water *ad libitum*. Measuring subject weight over time enabled careful monitoring of dehydration status. Subject weight was kept above 90% of adult, non-water restricted weight. The operant conditioning chamber was a transparent arena with three equally interspaced (distance of 10 cm) ports to record mouse responses and provide water rewards. The arena was adjacent to a linearized LCD screen (Viewsonic V3D245) that displayed visual stimuli. Mice licked the request port (center port) to display a visual stimulus on the monitor. Mice respond to the displayed stimulus by licking one of the response ports (left and right ports). Correct responses were rewarded with a small droplet of water (~ 10 µL) while incorrect responses were punished with a timeout (5-20 s). Auditory feedback consisting of beeps of various frequencies and white noise stimuli were also included to further indicate the nature of the responses: correct, incorrect, try again, etc. Throughout the training and testing process, we varied the reward size and the timeout duration to maintain high motivation in the subject.

### Task Sequence and Parameters of Stimuli for Behavior

Mice learned visual discrimination tasks over many weeks performing hundreds of trials a day and many thousand trials over the course of the experiment. High performance in the orientation discrimination tasks was achieved by taking the mice through a series of shaping trials. These trials trained naïve mice in using the operant chamber effectively, and in learning the structure of a self-directed two-alternateive forced choice (2AFC) trial before training them on orientation discrimination. We describe the shaping steps for two experiments relating to behavior: basic characterization of orientation tuning (Supplementary Table 1) and measuring integration times (Supplementary Table 2). Preliminary experiments determined the specific parameters used to probe behavior in the orientation discrimination task (data not shown). The screen subtended an angle of 100° X 65° (width X height) with respect to the subject. The spatial frequency of the gratings used for these experiments (0.08 cpd) maximized performance in most subjects. We used gratings tilted 45° to the vertical instead of vertical and horizontal gratings to prevent subtle variations in contrasts while rendering vertical vs horizontal gratings from influencing the behavior in an orientation independent fashion. While preliminary experiments used full-screen stimuli, all behavioral data shown in the paper used a circular aperture of diameter ~60°. This ensured that the spatial frequency of the stimulus at the edge of the aperture was no greater than 0.11 cpd. This spatial frequency was not so different as to change the overall performance of the animal and yet was high enough to allow multiple cycles of gratings within the aperture such that the mean luminance presented varied by less that 2% at different times during the trial (for drifting gratings) or for different trials (for flashed gratings). When different contrasts were shown on the screen, the stimuli presented were isoluminant with a mean luminance equal to the background luminance. While identifying the integration duration of subjects, we used flashed gratings with random spatial phases instead of drifting gratings because for the durations tested, the stimulus would have changed little and would have changed inconsistently from trial to trial. The randomized phases prevented the animal from using luminance information to perform the task – they had to use the overall angle of the grating presented to perform the task.

### Analysis of Behavior

We programmatically capture various facts about each trial performed by subjects. This allowed us to perform quantitative assessment of behavioral performance. In our tasks, the subjects have complete control over when trials are requested and when they respond to trial requests. For analysis, we excluded trials that were completed within 50 ms. Such fast responses would have required unattainable motor speeds and likely indicated water-clogged ports due to incomplete reward consumption. This condition excluded <0.1% of the trials. We further excluded trials that took >5 s. This was to ensure that we only counted trials where subjects were highly motivated to perform the task, excluding trials where subjects were distracted or in a low motivational state. The fraction of trials rejected due to large RTs did not exceed 3.05 % in any of our subjects and averaged 1.18 ± 0.91 %, (mean ± sd). We measured confidence intervals on the performance (# trials correct/# trials total) using the Clopper-Pearson method ^75^ and the significance of difference in binomial proportion using Agresti-Caffo statistics ^76^.

### Psychometric data fitting

In the experiments where we vary various features of the sensory input (contrast and duration), we model subject performance as a function of the strength of the stimulus:

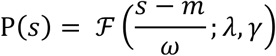

where ‘P’ is the psychophysical performance function,’ 𝓕’ is the logistic function, such that

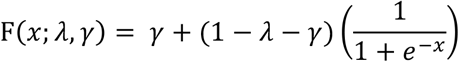

‘s’ is the strength of stimulus, ‘m’ is the stimulus strength at half-maximum performance, ‘ω’ is the width of the psychometric function (stimulus strengths where the psychometric function changes), ‘λ’ is the lapse rate and ‘γ’ is the guessing rate. In our analysis of contrast threshold, we define stimulus contrast at half-maximum performance as the threshold contrast (denoted as υ) and in our analysis of integration times, we define the stimulus duration at half-maximum performance as the threshold integration time (denoted as τ). Since all our behavioral data was obtained with a 2-AFC trial structure, we set γ to 0.5 in our analyses. Consistent with this assumption, when subjects were presented with trials having little or no information about the correct response (zero contrast stimulus or very low duration stimulus), subjects’ performance was no better than chance. We then identified those parameters (m,ω,λ) that best fit our data. To identify these parameters, we used constrained maximum likelihood techniques – techniques that maximizes the likelihood of the given data for some estimates of the unknown parameters, subject to constraints – to create point estimates. These analyses forced guessing rates to 0.5 and assumed that priors for the lapse rates (λ) followed a beta distribution with shape parameters (1.2, 12) while the priors for ‘m’ and ‘ω’ were flat through the stimulus range. We sampled from the posterior distribution using Markov-chain-Monte-Carlo techniques and used the 95-percentile range of the marginal of the posterior distribution of the fitted parameters as the confidence interval. These algorithms are based on previously described open-source methods (`psignifit`)^77^. We identify the integration time at the total time after which the performance of the animal does not change appreciably. This includes the time required to reach half-maximum performance (υ) and the time duration over which the performance changes from half-max to full max performance (ω/2). Thus, the total integration time is υ +ω/2.

### Animal variability and use of average subject

The perceptual thresholds measured varied from animal to animal. We fit psychometric tuning curves to the responses of individual animals and report the ranges of the fit parameters where applicable. When we report aggregate values, we used the values fit from the responses of an average subject. We simulated the average subject from the responses of all the animals in the population. However, different subjects performed different number of trials based on individual trial rates. Eight subjects were included in the contrast threshold estimation study where subjects performed between 2803 - 6078 trials averaging 4255 ± 1192 (mean ± sd). Six subjects were included in the visual integration study where subjects performed between 16098 - 54654 trials averaging 32051 ± 13896 (mean ± sd). To ensure equal weight for each animal in measuring aggregate fit parameters, we sampled the same number of trials from each subject randomly across stimulus conditions. We created the average subject by concatenating these sampled responses. To ensure that the sampling process did not bias estimates, we performed the resampling process 1000 times. We report the mean of the most likely estimates across these 1000 resamples for the average subject and plot the 95 percentile ranges for the same.

### Recording electrode implantation surgery for chronic probes

We used standard surgical techniques to implant Neuronexus probes in the superficial layer of the primary visual cortex. Adult mice were anesthetized under isoflurane (2.5% (v/v) for induction and 1.5% (v/v) for maintenance). After subjects were anesthetized, the fur from the top of the head was shaved and the mouse injected with atropine to minimize secretion (0.3 mg/kg), and dexamethasone to prevent inflammation (2.5 mg/kg). Subjects were then placed on a stereotaxic frame (Stoelting Co., Wood Dale, IL). The scalp over V1 was removed using surgical scissors and the skull dried. A small (<0.5 mm) craniotomy was made over the monocular region of V1 and Neuronexus probes (Neuronexus Inc, Ann Arbor, MI) were inserted (Poly2, Poly3 and A4×2-tet configurations) into the craniotomy. The open craniotomy was covered with silicone gel. A second adjacent craniotomy was made over the olfactory bulb and ground/reference screws were installed here. The exposed skull surface was then closed using dental cement. A custom designed head bar was also installed to enable headfixing for later experiments. Subjects were then removed from the stereotaxic frame and allowed to recuperate in a heated chamber and injected with buprenorphine for post-operative pain management (SQ, 0.3 mg/kg) All V1 recordings began at least 5 days post-surgery.

### Headcap surgery for acute recording preparation

The surgical procedure was identical to that of the chronic probe surgery in all respects except that no craniotomy was performed over V1. The skull surface was cleaned and the area over V1 was covered with a thin layer of cyanoacrylate-based glue and allowed to dry completely. The animal was allowed to recover for 5 days after surgery.

### Channelrhodopsin expression and fiber-optic cannula implantation

Channelrhodopsin-2 (ChR2) was targeted into Parvalbumin positive (PV+) fast spiking inter neurons within V1 through one of two ways. (1) PV-cre (JAX:0080609) subjects were injected with 200-400 nl of AAV2-Flex-ChR2-tdTomato(UPenn Vector core Cat#: AV-9-20297P) at various depths using a Nanoject II (Drummond Scientific, Broomall, PA) bilaterally over V1 and (2) PV-cre animals (JAX:0080609) were crossed with Flex-ChR2 (JAX:024109) and F1 progeny positive for both genes were selected for behavior. Fiber-optic cannulae (MFC_480/500-0.63_3mm_ZF1.25(G)_B60, Doric Lenses Inc., Quebec City, Quebec, Canada) were implanted into the open craniotomies over V1 such that the fiber terminus lay over the exposed skull. The open skull was covered with dental cement and the mouse removed from stereotaxic frame and set aside for recuperation. As a light stimulation negative control, two subjects underwent a sham injection. These subjects (PV-cre:0080609) were implanted with fiberoptic cannulae over V1 without viral injection.

### Optogenetic Manipulation During Behavior

Subjects implanted with fiberoptic cannulae were allowed to recuperate for 5 days after which they were introduced into the behavior chamber. We established baseline behavioral performance in the orientation discrimination task for ~ 1 week before attaching the animal to a fiber-optic light source. TTL pulses from the behavior computer controlled the LED light source (M470F3, ThorLabs, Newton, NJ, USA) through a LED Driver (LEDDB1, Thorlabs, Newton, NJ, USA). This light entered the behavior arena via an optical commutator (FRJ_1×1_FC_FC, Doric lenses, Quebec City, Canada) and then split into two through a 1×2 branching fiberoptic patch cord (BFP(2)_480/500/900-0.63_0.3m_FCM-2xZF1.25(F) Doric lenses, Quebec City, Canada) and mated with the cannulae with a plastic sleeve. Based on prior estimates of stimulus latency within V1^78^, we extended the LED light stimulation past the visual stimulus by 100 ms. Thus, LED lights started with stimulus onset and finished ~ 100ms after stimulus offset. We estimate the LED power to be ~9 mW at the fiber tip leading to a power density of ~12 mW/mm^2^.

### Stimulus presentation and electrophysiological recording from chronic probes

Subjects that underwent electrode implantation surgery were allowed to recuperate for 5 days. We acclimated subjects to being head fixed for 2 days before recording from V1 neurons. Subjects were head fixed by screwing the custom designed head bar to a mating bar. The mating bar was then attached to the recording rig with the mouse placed over a Styrofoam ball suspended in air to allow free range of motion ^79, 80^. Mice quickly adjust to being head fixed and begin running on the Styrofoam ball. Anecdotally, mice run >50% of the time (data not shown), but we did not measure running speed or pupil dilation during our recordings and could not correlate these features with spiking responses during our recording. While the mouse is head fixed, an LCD monitor (Viewsonic V3D245) is placed contralateral to the electrode implantation site. The LCD is the same distance away from the head fixed subject as it would be during behavior. The location of the monitor was adjusted to drive visual responses in the V1 neurons being recorded. The raw waveforms from V1 are buffered, filtered and digitized to a hard drive using the Open Ephys^81^ system. Concomitant to the physiological recording, we record synchronizing TTL pulses from the display computer to align spikes with the stimulus. Stimuli presented to the subject were similar in characteristics to the stimuli used to drive behavior except for a few characteristics. To maximize the number of neurons that were driven by stimuli, we chose to use full screen stimuli instead of through a circular window. An initial recording epoch collected the responses of subjects to gratings of different orientations (full contrast, 12 orientations, flashed for 500 ms or drifting for 2000 ms). The responses to these stimuli was used to characterize the orientation tuning of the neurons. Some sessions included responses to long duration stimuli (2000 ms) of full (100%) and low (15%) contrast drifting gratings (temporal frequency of 2Hz) tilted 45° from the vertical. After this characterization, we recorded responses of neurons to flashed gratings of short durations (50-200 ms). These included trials where the contrast presented was zero and no stimulus was shown on the screen. These trials measured background firing rates for the neuron.

### Stimulus presentation and electrophysiological recording from acute probes

Subjects were allowed to recover for 5 days post-surgery and were acclimated to the Styrofoam ball for at least 2 days before the recording. On the day of the recording, subjects were lightly anesthetized and a small craniotomy was performed over V1 and the exposed brain covered with silicone gel. The animal recovered from anesthesia for at least 2 hours before recording.

### Single unit identification

Raw neural data was filtered between 300 and 10000 Hz using a zero-phase digital filter. Open source libraries^82^ (`spikedetekt`) were used to detect putative spikes as significant voltage deviations (> 5 SD 2.5 SD from the mean voltage). Detected spikes were automatically clustered using an expectation maximization algorithm (`klusta`^82^) which modeled features (principal components of waveforms) of neurons as a mixture of gaussians. Clustered single units were manually verified using a visualization algorithm (`kwik-gui`). Over clustered units were combined based on the location of detected spike on the electrode, waveform shape, feature stability, and absence of refractory violations. Single units showed no refractory violation and were sufficiently separated from other units such that total false positive + false negative rates are less than 5%^83^. For each unit, we extracted the location within the brain calculated as the location of the electrode that had the largest mean waveform amplitude. This depth was used to categorize the unit as belonging to superficial Layer 2/3 (<400 μm) or deep Layer 5/6 (>450 μm). As we were recording with chronic electrodes, some units were — detected on the same electrodes across multiple days. These units were identified by looking for units with (1) the same waveform shape (>95% correlation across days) and (2) the same ISI distribution (>95% correlation across days) present at (3) the same depth across days. We found 190 V1 units that were present across multiple days. Duplicate units were removed such that only the responses of the units on the first day they were present was considered for future analysis.

### Analyzing V1 Responses

We synchronized spiking responses in V1 with stimulus presentation using TTL pulses. On each stimulus presentation, we extracted the total number of spikes in a time windows that started with the visual stimulus onset and extended to 500 ms after onset. We chose to analyze this time interval over a time interval cotemporaneous with stimulus presentation for three reasons. (1) V1 responses do not begin immediately after stimulus onset. Capacitive charging effects, and line delays cause response latency of ~50-100 ms^78, 84^. (2) V1 responses could sometimes extend far beyond the duration of the stimulus due to recurrent activity within the network^24^. (3) Analysis of various time intervals showed that a spike number code that included the chosen time interval (0-500 ms) maximized the average decoding performance across all neurons (data not shown), had close to the maximum number of consistent predictors of the stimulus (data not shown), and completely covered the stimulus in all conditions.

### Fitting Performance of Individual Neurons and for Populations of Neurons in a Session

Spike rates for each neuron was considered one at a time. We first excluded the firing rates for trials without stimuli (i.e. contrast=0). The firing rates for the remaining trials was randomly assigned to “Training” (70%) and “Testing” (30%). The data in the “Training” set was used to fit a logistic regression model (Matlab ^®^ function `mnrfit`) and logistic regression coefficients obtained for each neuron. This regression coefficient was used to test the model on the “Test” dataset (along with the no-stimulus trials) We repeated this process 100 times with a different subset of trials belonging to the “Training” and “Test” datasets. Neurons whose regression coefficients had the same sign for at least 70 out of the 100 attempts was considered a consistent predictor of the stimulus (i.e. they predict the stimulus as belonging to the same orientation no matter which trials are included in the fitting process). For fitting performance across populations, we used all the spike rates in a session and performed the regression in a similar fashion. The decoding performance of a single unit or of a population was the average performance across the 100 splits.

### Simulating Neural Subpopulations and Measuring Performance of Simulated Populations

For each of the neurons in our dataset, we tabulated the spike count responses separated by the stimulus conditions tested in our study (contrasts 0, 0.15,1 and durations from 50-200 ms). To create a virtual session from our dataset, we first chose a random subset from the overall population with replacement with each neuron being equally likely to be included in the sub-population. Responses for each neuron was then simulated from the corresponding response table making sure to only sample from the responses of that neuron for that stimulus condition. We used the previously calculated regression coefficients (calculated one neuron at a time) as the regression coefficient for the simulated neuron. The orientation of the stimulus was then predicted based on these independent regressors and compared against the input orientation. This was repeated 1000 times to provide an estimate of the performance of this population.

---

**Supplementary Figure 1:**
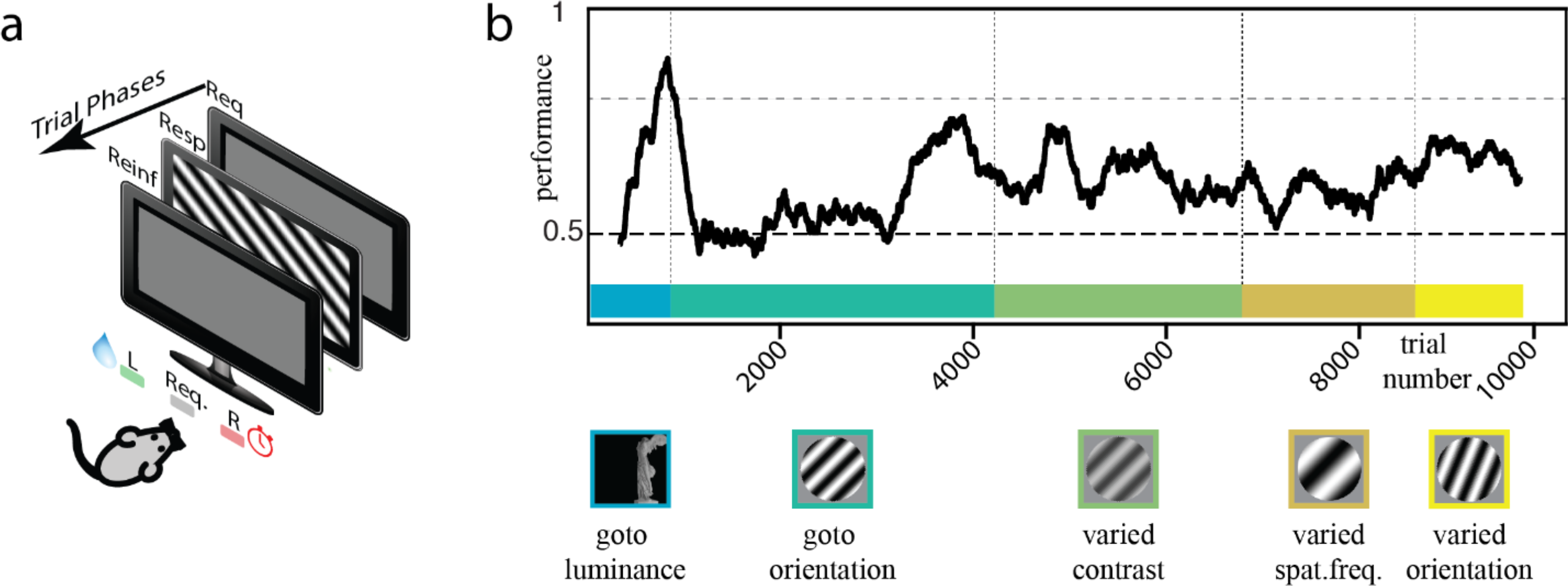
Mice are trained in visual discrimination in an automated operant conditioning chamber. **(A)** Schematic of trial structure performed by the subject. Subjects request trials during the Request phase (Req.) by licking the request port (center). Upon trial request, a visual stimulus is presented on the screen and the system waits for animal response (response phase; Resp.) The system then reinforces subject response in the reinforcement phase (Reinf.) by either providing water rewards (~10 ul) for correct responses or timeouts (5 – 20 s) for incorrect responses during which subjects cannot do any further trials. After the reinforcement phase, the system automatically returns to the request phase for the next trial. **(B)** Moving average of performance (black) of an example mouse through multiple steps (colored trial blocks) during the course of the experiment. Mice initially perform a go-towards-luminance task (blue). After reaching threshold performance, they are trained on a go-towards-orientation task (blue-green). After reaching threshold performance on this task, mice move into a series of modifications of the go-towards-orientation task where we varied various features of the stimulus(varied contrast, green; varied spatial frequency, orange; varied orientation, yellow) one at a time keeping all the others constant.

**Supplementary Figure 2:**
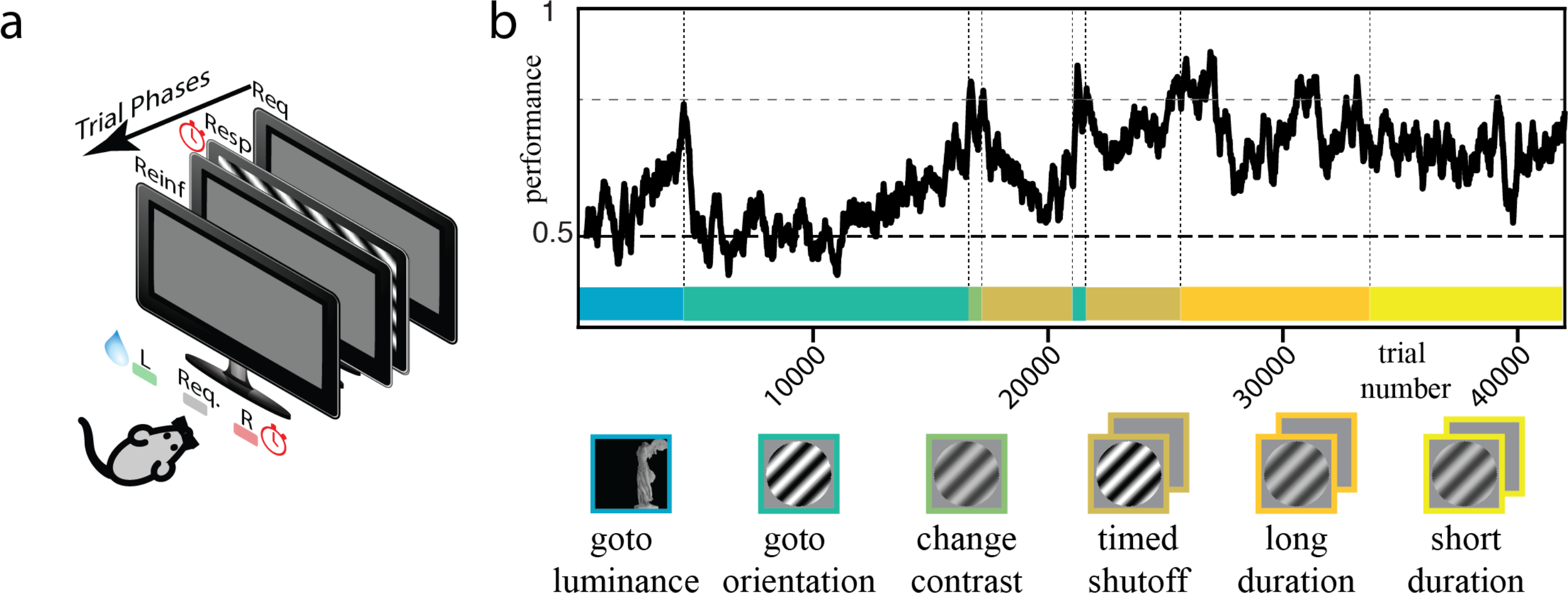
Mice went through a series of steps to measure integration times. **(A)** Schematic of trial structure performed by the subject. Subjects request trials during the Request phase (Req.) by licking the request port (center). Upon trial request, a visual stimulus is presented on the screen for a short duration before reverting to a grey screen and wait for subject response (response phase; Resp.) The system then reinforces subject response in the reinforcement phase (Reinf.) by either providing water rewards (~10 ul) for correct responses or timeouts (5 – 20 s) for incorrect responses during which subjects cannot do any further trials. After the reinforcement phase, the system automatically returns to the request phase for the next trial. **(B)** Moving average of performance (black) of an example mouse through multiple steps (colored epochs) during the course of the experiment. Mice initially perform a “go-towards-luminance” task (blue). After reaching threshold performance, they are trained on a “go-towards-orientation” task (blue-green). Subjects were then trained to perform OD on lower contrast trials “Change contrast” (green). In the timed shutoff step (brown) each trial request was followed by at most 1 second of high contrast stimulus following which the screen reverted back to gray screen. Subjects were forced to respond based on this limited information. After performing well on the times shutoff trials, subjects were shifted to trials where we varied the duration of the stimulus widely for durations between 120 ms – 2000 ms in the “long-duration” trials and between 16 ms – 250 ms in the “short-duration” trials. We manually moved the subject to an earlier task (~22000 trials) to improve motivation and prevent learned helplessness. The subject was shifted back to the timed-shutoff trials after ~1000 trials.

